# Detecting HTS Barcode Contamination

**DOI:** 10.1101/482166

**Authors:** Mallory A. Clark, Sara H. Stankiewicz, Vincent Barronette, Darrell O. Ricke

## Abstract

DNA barcoding enables multiple samples to be characterized in parallel with high throughput sequencing (HTS) experiments for cost efficiencies. Cross-contamination of DNA barcode reagents can result in the detection of HTS sequences for barcodes that were not originally added to a particular sample. Cross-contamination of data between multiplexed samples can also occur. Avoidance and detection of contaminated barcodes is relevant for DNA forensic samples analysis, accurate cancer diagnosis, clinical research applications, metagenomic analysis, etc. We present recommendations for the avoidance of contamination and a tool, TallyBarcodes, to aid in the detection of DNA barcode contamination.

## Introduction

DNA barcodes are short distinct segments of DNA that are used to identify one genetic sample from another. DNA barcodes enable scientists to run multiple samples in parallel, known as multiplexing[1]. Multiplexing reduces the costs and time of sequencing. However, DNA barcode reagents are susceptible to cross-contamination[2]. This contamination is easily seen when large counts for barcodes that were not used in a particular experiment are detected in the data. Finding non-random amounts of incorrect barcodes is evidence of cross-contamination in DNA barcode reagents[3]. Cross-contamination can also arise from laboratory experimental mistakes[4] [Bandelt & Salas 2009], during primer synthesis, multiple errors in HTS sequencing, or sample carryover from previous sequencing runs on the same instrument[5].

Sample misidentification was first observed by Kircher *et al.* [6] who recommended double indexing on the Illumina platform to mitigate the issue. Quail *et al.* [7] proposed SASI-Seq with sample assurance Spike-Ins to enable sample assurance. Wright & Vetsigian [8] reported read misassignments averaging at 0.24% on a Illumina HiSeq 2500; they propose quality filtering to alleviate sample cross-talk. Similar index misassignment rates between 0.06 and 0.21% are reported by[5, 9]. Sinha *et al.* [10] found that 5-10% of sequencing reads (signals) in a multiplexed pool of samples are mislabeled due to index switching on the Illumina HiSeq 4000 platform. Echoing Kircher *et al.* [6], MacConaill *et al.* [11] propose dual-indexed sequence adapters to eliminate index cross-talk. Esling *et al.* [2] also found that cross-contamination can occur between DNA barcodes and various primers and reagents. Integrated DNA Technologies (IDT)[12] reports 0.1% to 0.5% cross-contamination of HPLC-purified DNA oligos and 0.01% to 0.05% for Ultramer DNA oligos. To mitigate cross contamination risks in a laboratory, both Esling *et al.* [2] and Sinha *et al.* [10] recommend to construct bar code arrangements that do not overlap and to use uniformed lab handling and cleaning procedures[2, 10]. Some other recommended precautions are to label PCR replicates with one-time use tagged primers, prohibit single-tagging and double-tagging sample arrangements, and to not use long primer constructs for mixture samples [2].

## Materials & Methods

The software tool, TallyBarcodes, was developed to facilitate the detection of DNA barcode cross-contamination within experiments. TallyBarcodes enables the detection of contamination by quantifying the barcodes detected on the reads in the experimental data. TallyBarcodes is written in the Scala programming language.

## Ion Torrent HTS Sequencing

Thermo Fisher PGM, Proton, and S5 libraries were prepared and sequenced as described in Ricke *et al.* [13].

## Results

Two possible different types of sample misidentification from barcode cross-contamination are laboratory and n-1 series contamination. Both types of contamination have been detected in HTS experiment results. First, experimental results are consistent with laboratory contamination of barcode IX-08 with IX-16 during a specific experiment at a level of 0.0105 with standard deviation of 0.0016; the tube for barcode IX-16 is adjacent to IX-08 in a box of 16 barcodes with two rows of 8 tubes. Second, the n-1 series contamination patterns are characterized by trace contamination for a previous barcode reagent (n-1, n-2, etc.) in a sequentially numbered series for barcode reagents (n). Example n-1 series contamination patterns for different barcodes average 1.0e-4 for n-3 and n-2 to 1.5e-4 for n-1 and a maximum of 3.0e-4 to 5.0e-4 for n-1. Background random barcodes counts are less than 10 for HTS experiments averaging 100 million sequences.

## Discussion

While the source(s) of cross-contamination of DNA barcodes can remain unknown, instances of accidental cross-contamination of DNA barcode reagents has been observed in a research laboratory setting. The n-1 series contamination pattern that was observed is inconsistent with misread bases within the barcode sequence identified, assuming <1% base calling error rate[8]. Possible contributors of cross-contamination are splatter between tubes, incomplete purification (e.g., reuse of HPLC purification columns), etc.

Detection of individuals in complex DNA forensic mixtures, cancer cells, metagenomic community characterization, etc. are pushing the detection levels lower with advances in HTS technologies. Ricke *et al.* [13] can identify DNA contributors down to 1 in 400 concentration level. The n-1 series contamination levels of 3e-4 places a limit of 1 in 3,333, which is below current minor contributor detection level. The laboratory contamination of barcode IX-08 with IX-16 at 0.0105 is detectable at 1 in 95, which will prevent future combinations of these two barcodes in experiments until barcode IX-08 is replaced. A hypothetical combination of a normal individual sample with barcode IX-08 and a cancer sample or DNA mixture on barcode IX-16 could lead to incorrect conclusions from the results.

While certified laboratories have standard operating procedures (SOPs) to minimize cross-contamination between reagents, we present additional recommendations to reduce and detect contamination.

Recommendations:

- Consecutive experiments should use different barcodes
- Barcodes used together in one experiment should be combined with different barcodes in the next experiment to aid in detection of possible laboratory contamination
- Selected barcodes should be spaced to mitigate n-1 series contamination Double labeling of samples with two barcodes
- Clinical and DNA forensic samples should use special grade barcodes that have not been purified on equipment (e.g., HPLC columns) used to purify other barcodes
- Patient samples should not be multiplexed in the same experiments as cancer/tumor samples
- Forensic reference samples should be run in different experiments rather than mixture or crime scene samples
- Periodic screening of barcodes for possible contamination
- Replication of samples with different barcodes if contamination is suspected

The software tool, TallyBarcodes, is being provided as open source (https://github.com/doricke/TallyBarcodes) to facilitate characterization of HTS data.

DISTRIBUTION STATEMENT A. Approved for public release. Distribution is unlimited.

This material is based upon work supported under Air Force Contract No. FA8702-15-D-0001. Any opinions, findings, conclusions or recommendations expressed in this material are those of the author(s) and do not necessarily reflect the views of the U.S. Air Force.

